# *In vitro* reconstitution of calcium-dependent recruitment of the human ESCRT machinery in lysosomal membrane repair

**DOI:** 10.1101/2022.03.30.486420

**Authors:** Sankalp Shukla, Kevin P. Larsen, Chenxi Ou, Kevin Rose, James H. Hurley

## Abstract

The endosomal sorting complex required for transport (ESCRT) machinery has been shown to be centrally involved in repair of damage to both the plasma and lysosome membranes. ESCRT recruitment to sites of damage occurs on a fast time scale, and Ca^2+^ has been proposed to play a key signaling role in the process. Here, we show that the Ca^2+^-binding regulatory protein ALG-2 binds directly to negatively charged membranes in a Ca^2+^-dependent manner. Next, by monitoring the colocalization of ALIX with ALG-2 on negatively charged membranes, we show that ALG-2 recruits ALIX to the membrane. Furthermore, we show that ALIX recruitment to membrane orchestrates the downstream assembly of late-acting CHMP4B, CHMP3, CHMP2A subunits along with the AAA^+^ ATPase VPS4B. Finally, we show that ALG-2 can also recruit the ESCRT-III machinery to the membrane via the canonical ESCRT-I/II pathway. Our reconstitution experiments delineate the minimal sets of components needed to assemble the entire membrane repair machinery and open a new avenue for mechanistic understanding of endolysosomal membrane repair.

**Significance statement:** One of the ways by which protein aggregates can propagate and lead to progression of a neurodegenerative disease is by damaging the membrane that is destined to degrade the misfolded, aggregated protein. ESCRT machinery has been implicated in sealing these damaged membranes, and the nature of the membrane recruitment trigger signal for this machinery is a major open question. Here, we show *in vitro* that ALG-2 can bring ESCRT machinery to membranes in a Ca^2+^-dependent manner.

## Introduction

The endolysosomal system is susceptible to damage by a number of factors including protein aggregates, lysosomotropic compounds, reactive oxygen species, and lipid metabolites (1). Endosomal escape mediates the entry into the cytosol of many viruses (2), therapeutic agents (3, 4), and the transmission of prion-like molecular aggregates (5). Escape is counteracted by the protective effects of lysosomal membrane repair (6), which is carried out by the ESCRT membrane sealing machinery. The ESCRTs are a conserved membrane scission and sealing machinery consisting of about 30 proteins in humans (7, 8). In particular, ALIX, ESCRT-I and the ESCRT-III subunits CHMP6, CHMP2A, and CHMP2B have been implicated in cytoprotective lysosomal membrane repair events (6, 9, 10). ESCRTs are also involved in repair of the plasma membrane (11), where extracellular Ca^2+^ efflux has been shown to trigger the subsequent recruitment of ESCRT III machinery to the damaged plasma membrane site (12). The eventual closure of damage-induced holes in the plasma membrane is achieved by membrane budding and shedding vesicles towards the extracellular space by ESCRT III machinery (11).

Lysosomes have nearly a 5000-fold higher concentration of Ca^2+^ (~0.5 mM) compared to that in cytosol (~100 nM) (13, 14). Therefore, damage to the endolysosomal membrane locally increases Ca^2+^ concentration near the endolysosome. Increased local Ca^2+^ efflux into the cytosol has been proposed to be a trigger for the recruitment of endolysosomal membrane repair machinery. The Ca^2+^ binding protein ALG-2 co-accumulates with ALIX upon damage, and has been proposed to have an upstream role in the endolysosomal membrane repair sequence (9). ALG-2 contains five serially repetitive EF-hand structures and is the most conserved protein among the penta-EF-hand (PEF) family (15). Upon binding to Ca^2+^, ALG-2 undergoes conformational changes rendering it amenable to bind to proline rich proteins such as ALIX (16). Therefore, ALG-2 is a prominent candidate to trigger the recruitment of ESCRT III machinery, and therefore endolysosomal membrane repair, in response to increased local Ca^2+^ concentration around a damaged endolysome. However, whether ALG-2 is sufficient to trigger the membrane recruitment of repair machinery at the site of damage is unclear (17).

Here, we investigated the mechanism directly through *in vitro* reconstitution in a completely defined system of purified proteins and synthetic lipids. *In vitro* reconstitution is a powerful tool to determine the sufficiency of a biochemical factors, which we applied here to probe whether ALG-2 and. We used a giant unilamellar vesicle (GUV) reconstitution system. We first showed that ALG-2 can be recruited to the negatively charged membranes without the need for an upstream adaptor protein. Next, by using a Ca^2+^ binding deficient mutant of ALG-2, we confirmed that ALG-2 membrane recruitment is mediated by Ca^2+^. Subsequently, by monitoring the colocalization of ALIX with ALG-2 on the negatively charged membrane, we attest to the upstream role of ALG-2 in bringing ALIX to the membrane. Furthermore, we show the complete downstream recruitment of human ESCRT III machinery vis-à-vis CHMP4B, CHMP2A, CHMP3 along with the AAA^+^ ATPase VPS4B to the negatively charged membranes in an ALG-2 and Ca^2+^ dependent manner. Additionally, we demonstrate that ALG-2 also recruits the ESCRT-III machinery via the canonical ESCRT-I, ESCRT-II pathway. Finally, we validate that endolysosomal membrane damage leads to colocalization of ALG-2, ALIX, and ESCRT-I *in vivo*.

## RESULTS

### ALG-2 binds to negatively charged membranes in a Ca^2+^-dependent manner

We set out to decipher the endolysosomal membrane repair machinery upstream of ESCRT III recruitment at the site of endolysosome membrane damage. To do this, we used *in vitro* GUV reconstitution experiments to sequentially test the membrane binding ability of the hypothesized endolysosomal membrane repair machinery. We incubated GUVs containing 30% DOPS (along with 69.5% DOPC and 0.5% Atto 647 N DOPE) with ALG-2 (Atto 488; 200 nM) for 15 minutes in reaction buffer. On imaging, we observed ALG-2 puncta on the GUVs (Fig. 1A), suggesting that ALG-2 can bind to negatively charged membranes without the need for an upstream membrane anchor. Surprisingly, membrane recruitment of ALG-2 occurred in the absence of externally added Ca^2+^. Furthermore, external addition of Ca^2+^ to a concentration of 100 μM did not affect the binding of ALG-2 to membranes.

**Figure 1.**
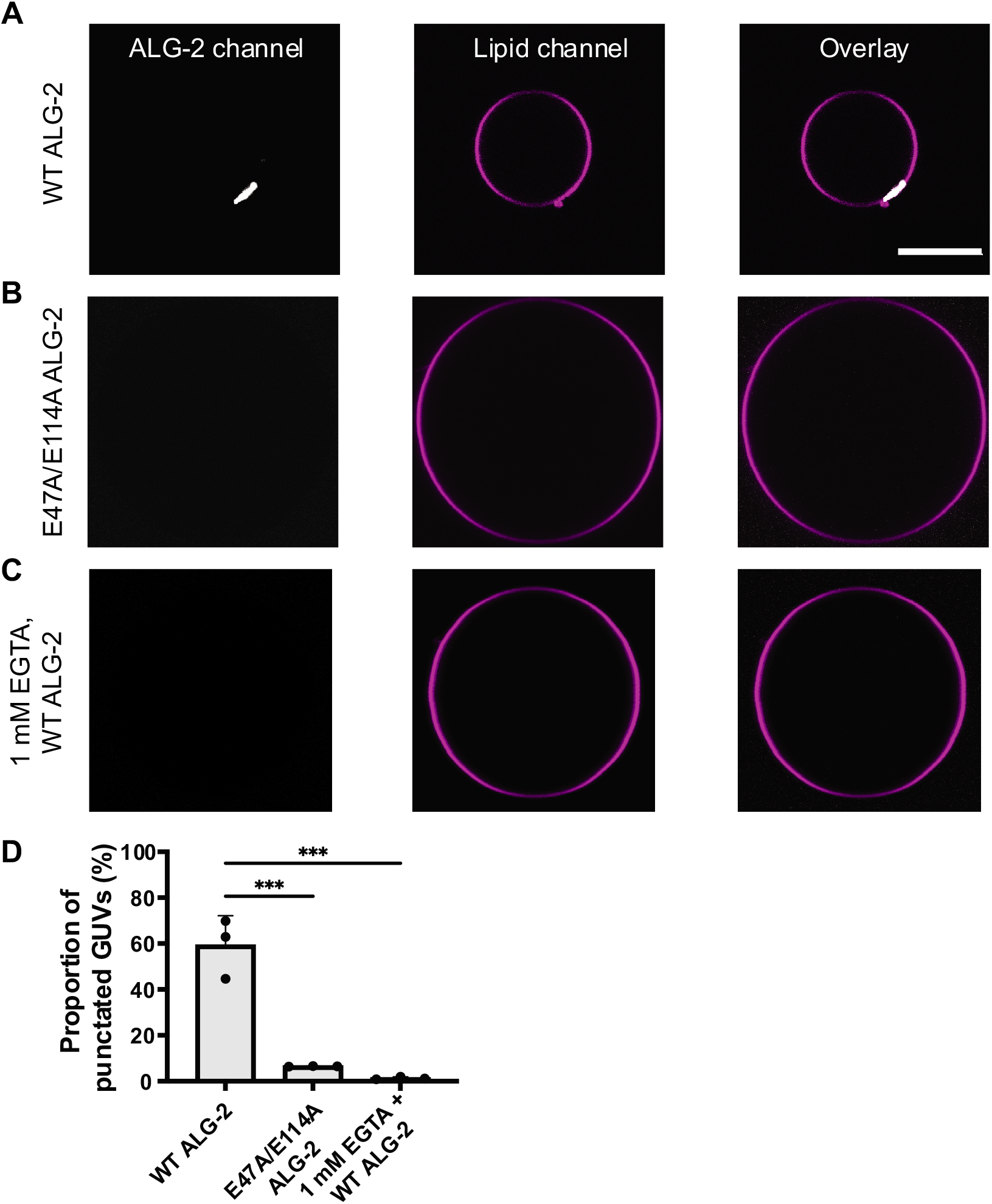
Membrane recruitment of ALG-2 is Ca^2+^-dependent and electrostatic in nature. The GUVs were prepared using the PVA-gel based protocol described in the Materials and Methods. ALG-2 was fluorescently labeled using Atto 488 maleimide by introducing a solvent accessible mutation, A78C in WT ALG-2. a) Atto 488 labeled ALG-2 is recruited to the periphery of the 30% DOPS containing GUVs. The periphery of a GUV was determined by using the fluorescence of the lipid dye (Atto 647 N – DOPE) added in trace amounts (0.5%) to the lipid mixture used for GUV preparation. b) shows the absence of ALG-2 recruitment to GUV periphery on replacing the WT ALG-2 to the Ca^2+^ binding deficient mutant of ALG-2 (E47A/E114A). c) Using an ALG-2 stock incubated with 1 mM EGTA overnight also abrogated the membrane binding of ALG-2. The absence of ALG-2 puncta on GUVs confirm that the membrane recruitment of ALG-2 is calcium dependent. d) The proportion of GUVs that had at least one ALG-2 puncta on their periphery were plotted for fluorescently labeled WT ALG2, E47A/E114A ALG2, and 1 mM EGTA incubated ALG-2. Data points are mean ± SD. p ≤ 0.003 (***). Scale bar is 10 μm.

We hypothesized that ALG-2 was purified in the Ca^2+^-bound activated form and therefore was already in its activated Ca^2+^ bound form. To test our hypothesis, we purified a Ca^2+^ binding deficient mutant (E47A/E114A) (15) of ALG-2 and performed the same membrane binding experiment. We found that the Ca^2+^-binding deficient mutant (E47A/E114A) of ALG-2 did not bind 30% DOPS containing membranes (Fig. 1B). Moreover, addition of 100 μM CaCl_2_ to ALG-2^E47A/E114A^ had no effect on its membrane non-binding behavior. This confirmed that at a given negative charge density the membrane binding ability of ALG-2 is Ca^2+^-dependent. Additionally, we incubated a purified stock of ALG-2 (50 μM) with 1 mM EGTA (pH 7.4) overnight. On using the EGTA incubated ALG-2 stock to perform membrane binding experiments, we did not observe ALG-2 puncta on the membrane (Fig. 1C). Together with the results for ALG-2^E47A/E114A^, we confirmed that Ca^2+^ is the trigger for recruiting ALG-2 to the negatively charged membranes.

Earlier work on plasma membrane repair machinery has proposed ALG-2 to be an important component. In the reconstitution experiments performed by Sonder et al., they reported that ALG-2 does not bind 10 mol% DOPS membranes on its own. The authors reported that the Ca^2+^-dependent binding of Annexin 7 is necessary to bring ALG-2 to negatively charged membranes (17). However, the amount of negatively charged lipids in membranes used in their study was 10 mol%, which also does not support strong membrane binding in our experiments. Since the membrane binding ability of ALG-2 is at least in part electrostatic in nature, therefore, this could explain why Sonder et al. did not observe membrane binding of ALG-2.

### ALG-2 is necessary for membrane recruitment of ALIX

ALIX is an ESCRT accessory protein that forms an alternative pathway to that of ESCRT I-II for the recruitment and activation of ESCRT-III in humans (18). ALIX consists of a BRO1 domain, a V domain and a proline rich domain (PRD), and functions as a homodimer (19). Earlier work has shown that ALIX in resting state exists in an autoinhibited conformation (20) and needs an upstream factor for its activation. The upstream binding of ALG-2 to the PRD of ALIX in a Ca^2+^-dependent manner has been shown to release the autoinhibition of ALIX and make the Bro1 domain accessible to bind downstream ESCRT III machinery (21). With this understanding, we incubated ALG-2 (Atto 488; 200 nM) with ALIX (Cy3; 100 nM) in the reaction buffer for 15-min before mixing them with 30% DOPS GUVs. We found that ALIX is recruited to the negatively charged membranes (30% DOPS) only when ALG-2 is present (Fig. 2A). In the absence of ALG-2, we did not observe membrane binding of ALIX (Fig. 2B). External addition of CaCl_2_ to the ALIX-only sample did not promote membrane binding of ALIX.

**Figure 2.**
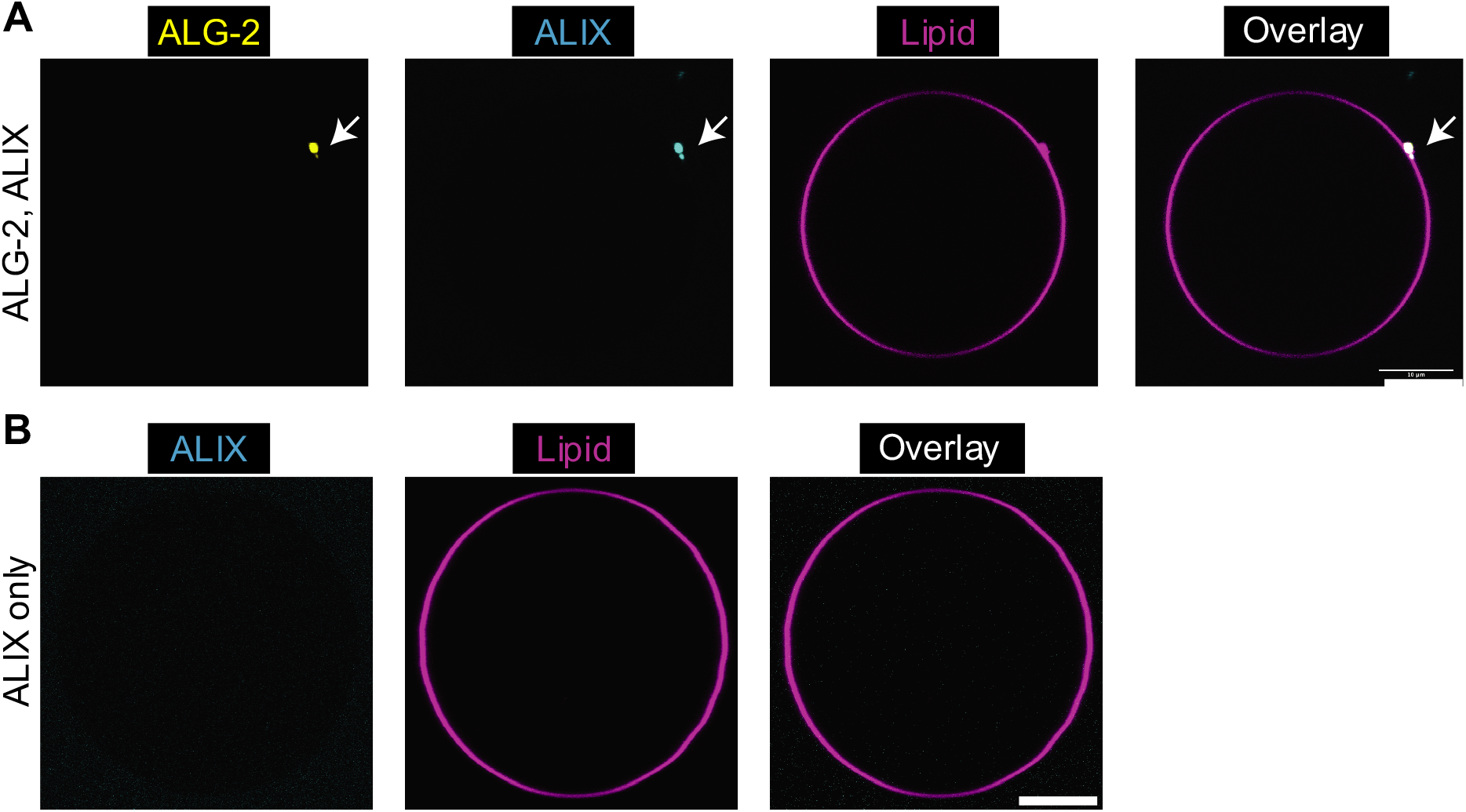
ALG-2 activates and co-recruits ALIX to membranes. a) GUVs containing 30% DOPS are incubated with fluorescently labeled ALG-2 (Atto 488; 200 nM) and ALIX (Cy3; 100 nM). We observe a clear colocalization between the membrane recruited ALG-2 and ALIX. b) In the absence of ALG-2, no ALIX puncta were observed on the membranes. Scale bar is 5 μm.

Additionally, ALIX was not recruited to membrane by the Ca^2+^-binding deficient mutant of ALG-2. The reason for non-observance of ALIX puncta on the membrane on replacing WT ALG-2 with Ca^2+^-binding deficient mutant of ALG-2 could be two-fold. First, the interaction between ALIX and ALG-2 has been shown to be mediated by Ca^2+^ (22). Second, based on our earlier observation, ALG-2 membrane interaction is Ca^2+^-mediated. Therefore, even if there is interaction between ALIX and Ca^2+^-binding deficient mutant of ALG-2, ALIX will not be recruited to membranes. Furthermore, we checked that ALG-2 dependent ALIX recruitment to membrane (and conversely, membrane non-binding of ALIX by Ca^2+^-binding deficient mutant of ALG-2) existed for labelled as well as dark ALG-2 suggesting that fluorophore labeling of ALG-2 did not interfere with the recruitment of endolysosomal membrane repair machinery. Therefore, for subsequent experiments, we used dark ALG-2 in conjunction with labelled downstream proteins of the endolysosomal membrane repair machinery to prevent bleed through between imaging channels.

### The downstream ESCRT-III machinery is recruited to ALG-2/ALIX puncta

ALIX has been shown to recruit the downstream ESCRT III machinery to membrane upon an upstream cue (18). In the cytosol, ESCRT-III proteins exist in an auto-inhibitory state (23). The C-terminal autoinhibition needs to be released for the ESCRT-III proteins to be functional (24, 25). Specifically, for CHMP4B, the Bro1 domain of ALIX interacts with the alpha 6 of CHMP4B to relieve its autoinhibition and convert it to an open conformation. CHMP4B (and other ESCRT-III proteins) electrostatically bind to negatively charged membranes through their electropositive amino-terminal residues as well as through insertion of their N-terminal amphipathic helix (26).

CHMP4B, CHMP2A, and CHMP3 along with VPS4B are the human counterparts of the minimal ESCRT III machinery needed for membrane scission (27). We found that under our experimental conditions (reaction buffer and 30% DOPS GUVs), CHMP4B can bind to membranes without any upstream factor at a concentration of 50 nM. At this concentration, CHMP4B coats the entire membrane surface within minutes after incubation with GUVs. We speculate that membrane plays a role in the release of auto-inhibition of CHMP4B at this concentration, an effect which could be allosteric in nature.

Consistent with earlier experiments (18), we found that CHMP4B is necessary to recruit the downstream heteropolymers of CHMP2A and CHMP3. Addition of 100 nM CHMP2A and CHMP3 each to 50 nM CHMP4B resulted in the recruitment of CHMP2A/3 heteropolymers to GUVs. Neither CHMP2A nor CHMP3 were recruited to 30% DOPS GUVs at a concentration of 100 nM each in the absence of CHMP4B. Therefore, at a concentration of CHMP4B where it can bind to membranes on its own, the downstream ESCRT III machinery also gets recruited to the membrane.

However, physiologically, ESCRT-III proteins are cytosolic in their resting state and are recruited to membranes only in response to an upstream cue (28). Accordingly, to mimic the physiological machinery, we decreased the concentration of CHMP4B to avoid its self-recruitment to membranes. We found that at a concentration of 10 nM, CHMP4B no longer gets recruited to 30% DOPS GUVs by itself. At this concentration of CHMP4B, addition of ALG-2 (200 nM) and ALIX (100 nM), resulted in the recruitment of CHMP4B to the membrane. The membrane recruited CHMP4B colocalized with ALG-2/ALIX puncta (Fig. 3A, row 1; 3B). Additionally, omission of ALIX prevented recruitment of CHMP4B to membrane (Fig. 3A, row 2; 3B).

**Figure 3.**
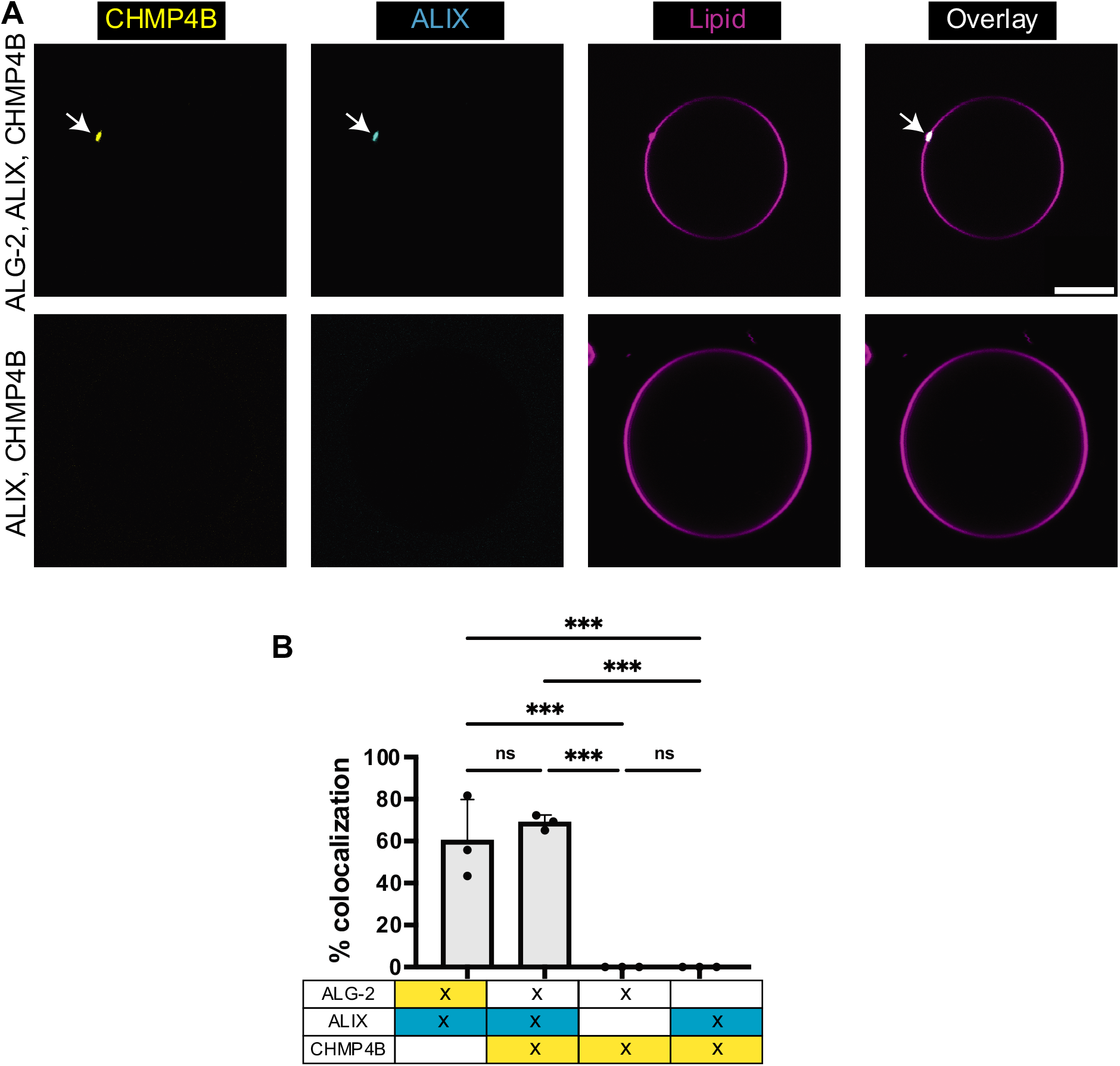
CHMP4B is recruited to membranes by ALIX. a) Addition of ALG-2 (dark; 200 nM) and ALIX (Cy3; 100 nM) along with CHMP4B (Atto 488; 10 nM) to 30% DOPS containing GUVs results in colocalized puncta of ALIX and CHMP4B. In the absence of ALG-2, there were no observable membrane recruitment of ALIX and CHMP4B onto the membrane. b) shows the histogram depicting the colocalization % of the puncta in two different fluorescently labeled protein channels. Data points are mean ± SD. p – 0.0003 (***). Scale bar is 10 μm.

### Complete reconstitution of ESCRT III machinery in response to Ca^2+^ binding by ALG-2

It has been shown that AAA^+^ ATPase VPS4 is necessary for membrane scission by virtue of disassembling ESCRT-III polymers on membrane necks (29). The established recycling function of VPS4 depends on the interaction of the MIT domain in VPS4 with MIT-interacting motifs (MIMs) in ESCRT-III subunits (30, 31). Therefore, membrane recruited upstream ESCRT III proteins should also recruit VPS4B. To test this in our experiments, we added 100 nM VPS4B and saw its colocalization with 50 nM CHMP4B, 100 nM CHMP2A, and 100 nM CHMP3 on 30% DOPS membranes. In the absence of ESCRT III proteins, VPS4B did not get recruited to membranes even after increasing its concentration to 500 nM.

For the complete reconstitution of the endolysosomal membrane repair machinery, we used fluorescently labeled ALIX (Cy3; 100 nM), CHMP4B (Atto 488; 10 nM), and VPS4B (LD 655; 100 nM). All other proteins involved in the reconstitution machinery, namely, ALG-2 (200 nM), CHMP2A (100 nM), and CHMP3 (100 nM) were unlabeled. All the abovementioned proteins were mixed in the reaction buffer at the aforementioned concentrations and incubated for 30 mins before adding GUVs. The imaging was started 15 mins after GUV addition. We found that all the three labeled proteins were colocalized on the GUV surface (Fig. 4A). Additionally, omission of either CHMP4B or CHMP2A/3 resulted in the abrogation of membrane recruitment of VPS4B (Fig. 4B). The colocalization statistics of these experiments are shown in Fig. 4C.

**Figure 4.**
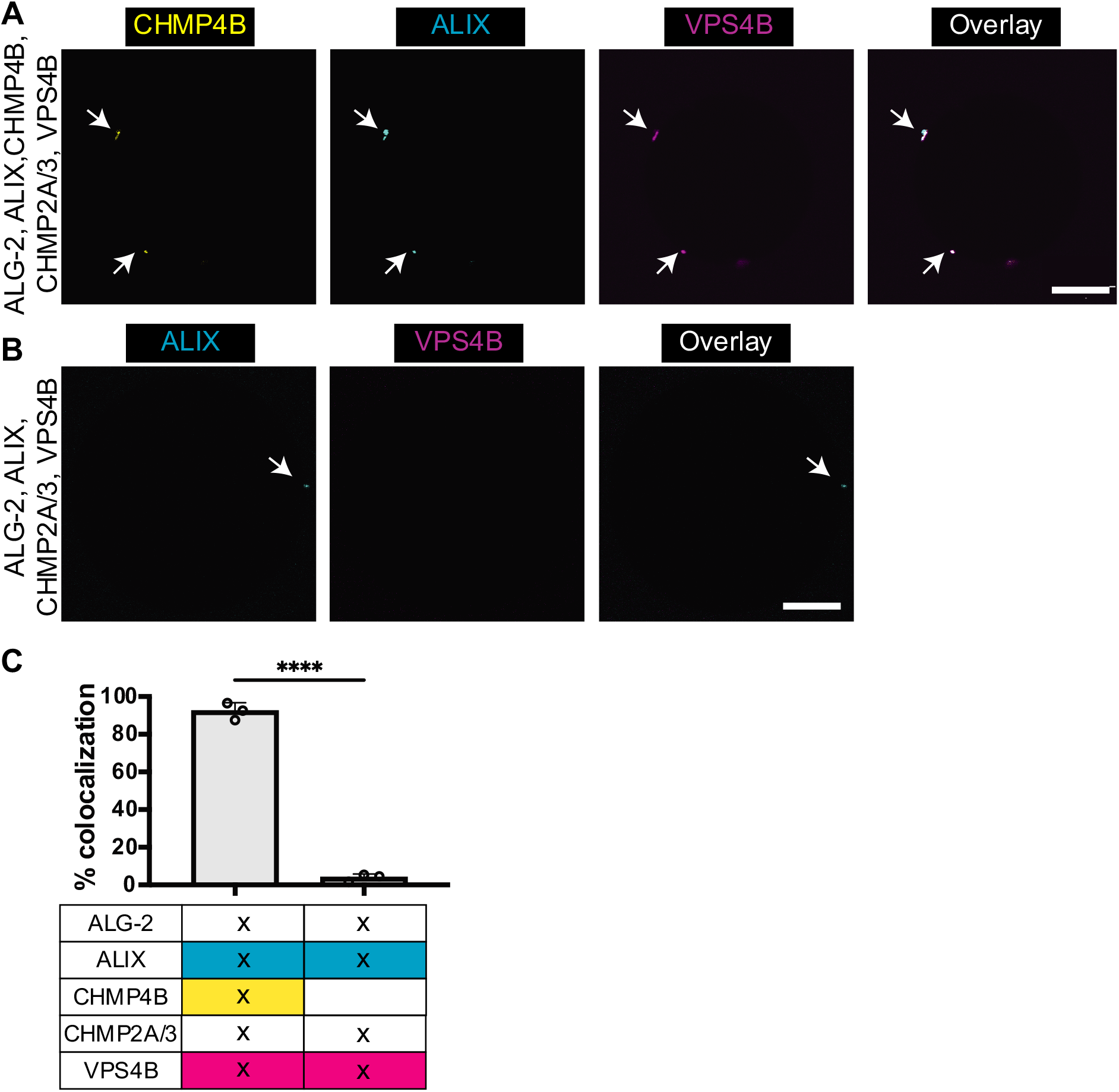
Complete reconstitution of Ca^2+^-triggered endolysosomal membrane repair machinery. A) Fluorescently labeled ALIX (Cy3; 100 nM), CHMP4B (Atto 488; 10 nM), and VPS4B (lumidyne 655; 100 nM) along with ALG-2 (dark; 200 nM), CHMP2A (dark; 100 nM), and CHMP3 (dark; 100 nM) are mixed with 30% DOPS containing GUVs and imaged. The observance of colocalized puncta of the labeled proteins on the periphery of GUVs confirms the recruitment of the entire endolysosomal membrane repair machinery. B) In the absence of CHMP4B, labeled VPS4B was not recruited to the ALIX puncta on the membrane. C) shows the histogram depicting the colocalization % of the puncta in two different fluorescently labeled protein channels. Data points are mean ± SD. p<0.0001 (****). Scale bar is 10 μm.

Above, we showed that ALIX is recruited to membranes only when ALG-2 is bound to membranes. We have also shown that VPS4B is recruited to 30 mol% negatively charged membrane only when CHMP2A and CHMP3 are present. Therefore, recruitment of VPS4B to membranes is a confirmation that CHMP2A and CHMP3 are also colocalized at the membrane. In conclusion, we demonstrate the complete reconstitution of the ALIX mediated endolysosomal membrane repair machinery, starting from the Ca^2+^-activated ALG-2 to the AAA^+^ ATPase VPS4B.

### ALG-2 recruits ESCRT-I complex to negatively charged membranes

ESCRT-I, ESCRT-II, and CHMP6 form the canonical pathway for the recruitment of ESCRT-III machinery (18). Human ESCRT-I is a soluble hetero-tetramer consisting of TSG101, VPS28, VPS37(A-D) and MVB12 (A, B). Earlier mutation-based experiments have shown the interaction between the proline rich region of the TSG101 subunit of ESCRT-I with ALG-2 in a Ca^2+^-dependent manner (32). Also, it has been observed *in vivo* that membrane repair is completely stalled only on co-depletion of ALIX and TSG101 and not on individual depletion of either of these proteins (9). When depleted individually, loss of TSG101, but not ALIX, was shown to reduce cell viability (10). These data implied a central role for ESCRT-I and motivated us to ask the question whether ALG-2 could be the upstream connecting link in recruiting ESCRT III machinery for membrane repair via ESCRT-I.

To do this, we incubated fluorescently labeled ALG-2 (Atto 488; 200 nM) with ESCRT-I complex (Cy3; 50 nM) in the reaction buffer for 15-min. We then added 30% DOPS GUVs to this mixture and imaged them after a 15-minute incubation. We found that the ESCRT-I complex is colocalized with the ALG-2 puncta on the membrane (Fig. 5B). In our experiments, we did not see membrane recruitment of ESCRT-I in the absence of ALG-2 (Fig. 5C).

**Figure 5.**
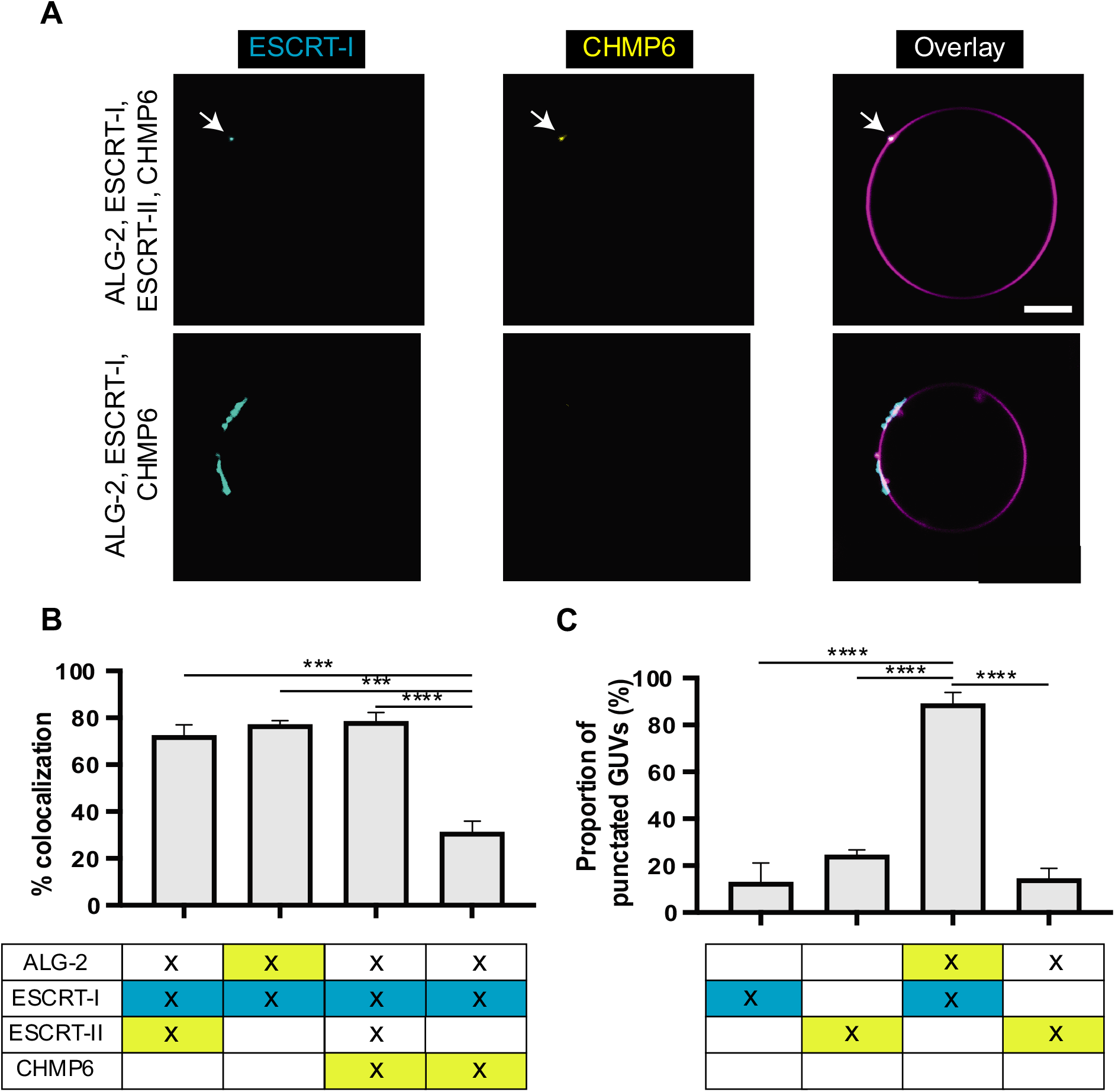
ALG-2 mediated canonical ESCRT III recruitment pathway. a) (Row 1) GUVs containing 30% DOPS are incubated with ALG-2 (dark; 200 nM), ESCRT-I (Cy3; 50 nM), ESCRT-II (dark; 200 nM), and CHMP6 (Atto 488; 400 nM). Imaging post 15-minute of incubation resulted in ESCRT-I, CHMP6 colocalized puncta on the surface of GUVs. This confirmed that the ALG-2 recruits the downstream cascade of ESCRT III machinery recruitment via ESCRT-I and ESCRT-II. (Row 2) On omitting ESCRT-II, the CHMP6 puncta on the membrane were mostly missing. The CHMP6 puncta that were occasionally observed rarely coincided with the ESCRT-I puncta. b) depicts histograms for the colocalization % of the puncta observed for membrane recruited membranes. c) shows the proportion of GUVs that had atleast one puncta for the conditions specified underneath each histogram. Data points are mean ± SD. p = 0.0002 (***), p<0.0001 (****). Scale bar is 10 μm.

Next, we asked whether we could reconstitute the canonical pathway of ESCRT III recruitment downstream of ESCRT-I. Human ESCRT-II is a Y-shaped tetrameric complex comprised of two EAP20 subunits and one copy each of EAP30 and EAP45 (33–35). It has been shown that the VPS28 subunit of the ESCRT-I complex interacts with the EAP45 subunit of the ESCRT-II complex (36). Therefore, we incubated fluorescently labeled ESCRT-II (Atto 488; 200 nM) complex with ESCRT-I (Cy3; 50 nM) complex and ALG-2 (dark; 200 nM) for 15-minutes. Thereafter, GUVs were added and imaged after 15 min of incubation. We found that both ESCRT-I and ESCRT-II colocalize on the GUVs (Fig. 5B). In contrast, we did not observe membrane binding of ESCRT-II in the absence of either ALG-2 or ESCRT-I (Fig. 5C).

Furthermore, it is known that the EAP 20 subunit of ESCRT-II binds to the N-terminal half of CHMP6 (33, 37). Therefore, we next added CHMP6 to the mix. To confirm that the entire pathway starting from ALG-2 to CHMP6 is being sequentially recruited onto the membranes, we used ALG-2 (dark; 200 nM), 50 nM Cy3 labeled ESCRT-I (Cy3; 50 nM), ESCRT-II (dark; 200 nM), and CHMP6 (Atto 488; 400 nM). We observed colocalized puncta of CHMP6 with ESCRT-I (Fig. 5B). Omitting either ESCRT-I or ESCRT-II abrogated the membrane recruitment of CHMP6 (Fig. 5C). Together, this confirmed that the canonical pathway of ESCRT-III recruitment to membrane is orchestrated by ALG-2.

### ALG-2, TSG101, and ALIX are co-recruited to the sites of lysosomal damage

Earlier cell-based experiments have shown co-accumulation of ALG-2 with ALIX upon endolysosomal membrane damage (9), as well as co-accumulation of ESCRT-I subunit TSG101 with the ESCRT-II subunit EAP30 (10). The association of ESCRT-I and ALG-2 has not been directly explored, however. Through our *in vitro* reconstitution experiments, we found ALG-2 to be a viable upstream candidate to recruit the ESCRT III machinery to the negatively charged membranes via the ALIX and the ESCRT-I/II pathway. Therefore, to validate our findings, we treated U2OS cells with the membrane-permeant lysosomotropic agent L-Leucyl-L-Leucine methyl ester (LLOME) to induce rapid nanoscale ruptures of endolysosomal membranes (38, 39). Once in the lumen of the lysosomal membrane, LLOME becomes acidified and condenses into polymers that reparably damage the lysosomal compartment. Lysosomal membrane ruptures result in the luminal accumulation of otherwise galectin 3 (Gal3). Therefore, damaged lysosomes can be distinguished from other acidic organelles by virtue of a detectable Gal3 signal. We confirmed that acute LLOME treatment (15-minute exposure) resulted in a dramatic shift of diffusely cytoplasmic Gal3 (Fig. 6A, upper panel) to punctate structures resembling membranous organelles (Fig. 6A, lower panel). On these damaged lysosomes, immunostaining confirmed the accumulation of ALG-2 and ALIX as previously shown. Additionally, in tune with our GUV-based findings, we observed colocalization of ESCRT-I component TSG101 with ALG-2 at these sites which also coincided with ALIX (Fig. 6A, lower panel). The statistically significant increase in the coincidence area of ESCRT-I, ALIX, ALG-2, and Gal3 upon LLOME treatment (Fig. 6B) confirms corecruitment of both ALIX and ESCRT-I with ALG-2 on the damaged endolysosomes.

**Figure 6.**
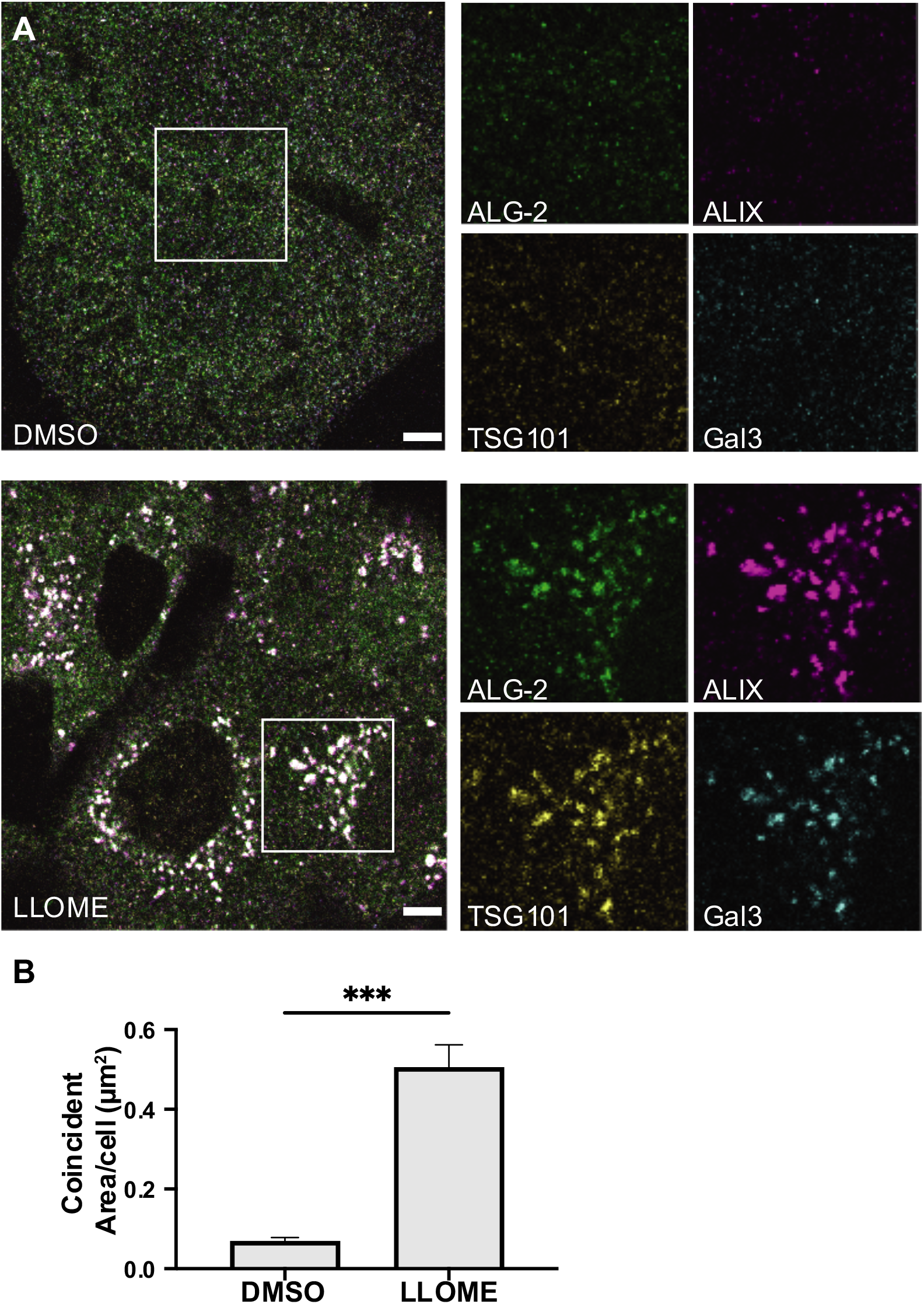
ALIX and the ESCRT-I recruitment factor TSG101 are co-recruited with ALG-2 to damaged lysosomes. (A) U2OS cells were treated with either DMSO (upper panel) or LLOME (lower panel) for 15 minutes and were then immunostained for ALG-2 (green), ALIX (magenta), TSG101 (yellow), and Gal3 (cyan). Relatively large colocalized puncta of ALG-2, ALIX, and TSG101 alongwith Gal3 were observed after LLOME treatment. The regions encapsulated within the white boxes have been enlarged and the individual protein channels are shown on the right. Images are representative of 5 independent replicates. (B) The area of overlapping compartments increased dramatically upon LLOME treatment compared to the DMSO control (n=75 cells for DMSO, 64 cells for LLOME). Data points are mean ± SD. Scale bar is 5 μm.

## Discussion

In this study, through our membrane reconstitution experiments with human ESCRT III proteins, we substantiated the concept that Ca^2+^ and ALG-2 are a major trigger for rapid recruitment of ALIX and ESCRT-I to sites of damaged endolysosomal membrane. A major and unexpected conclusion drawn from our data is that membrane recruitment of ALG-2 occurs in a Ca^2+^-dependent manner without the need for other proteins. The membrane-recruited ALG-2 forms puncta which could imply formation of a higher order assembly of ALG-2 on membranes, a topic calling for further exploration. Our observations are consistent with the model proposed by Scheffer *et al*. for plasma membrane repair (12), suggesting close parallels between the processes at the plasma membrane and lysosomes. Subsequently, we showed the downstream recruitment of the entire ESCRT-III machinery to the membranes with Ca^2+^ bound ALG-2 as the trigger. Most analysis of the ALG-2 pathway in membrane repair has focused on ALIX (9, 11, 12, 40), yet TSG101 is the more important contributor to maintaining cell viability under lysosomal damage conditions (10). Here, we demonstrated that ALG-2 can also recruit ESCRT-I to the membranes in a Ca^2+^-dependent manner. Subsequently, this leads to the recruitment of downstream ESCRT-II complex and CHMP6 which is part of the ESCRT-III machinery. The downstream recruitment of the ESCRT-III machinery in response to Ca^2+^ efflux from the damaged endolysosome could potentially lead to the endolysosomal membrane repair through membrane constriction and scission facilitated by a coherent action of ESCRT-III proteins and AAA^+^ATPase VPS4B (Fig. 7).

**Figure 7.**
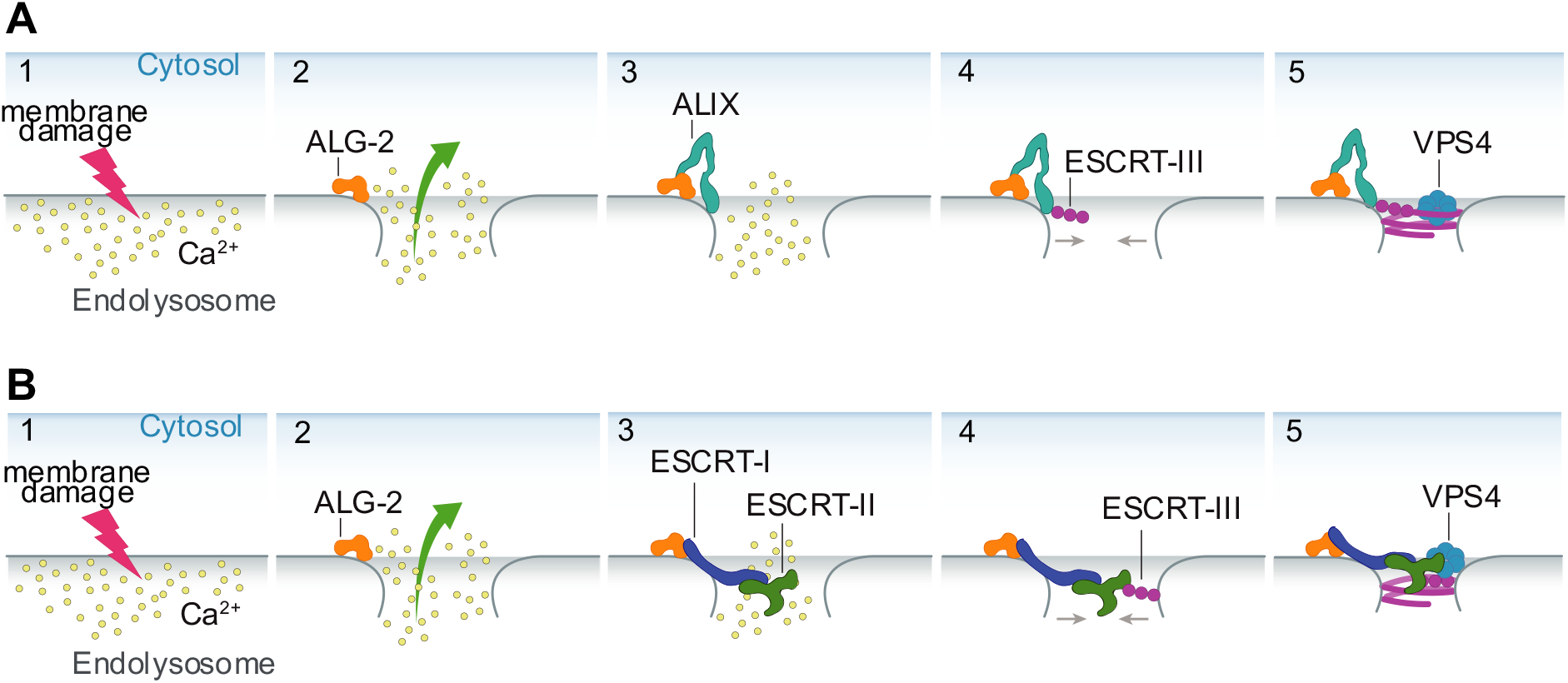
Schematic depicting the Ca^2+^-triggered endolysosomal membrane repair model. A cartoon representation of Ca^2+^ triggered endolysosomal membrane repair by ESCRT-III machinery mediated by ALIX (panel A) or ESCRT-I/II (panel B). The endolysosomal membrane damage leads to a Ca^2+^ efflux from the endolysosome into the cytoplasm (A, B; 1) which leads to the membrane recruitment of Ca^2+^ chelator, ALG-2 (A, B; 2). The membrane recruited ALG-2 leads to a downstream recruitment of ESCRT-III proteins via two parallel pathways: the ALIX pathway (A; 3, 4) and the canonical ESCRT-I/II pathway (B; 3, 4). The assembled ESCRT-III machinery forms a higher order structure that can bring the open ends of the damaged membrane together (A, B; 5). Finally, disassembly of ESCRT III filaments by AAA^+^ ATPase VPS4B potentially leads to membrane closure and repair.

Recent cell based endolysosomal membrane damage studies have shown a fast recruitment (within minutes) of ESCRT III machinery to the membrane damage sites compared to the delayed localization of the initiators of autophagic machinery (9, 10). The arrival of ESCRT III machinery in response to sites of membrane damage coincided with the restoration of membrane integrity and function (9, 10). While the recruitment of ESCRT III machinery has been conclusively shown to respond to membrane damage, the specific signals following membrane damage, which trigger ESCRT protein recruitment and the mechanistic roles of ESCRT machinery in membrane repair have remained elusive. Our study fills this gap in understanding by showing that ALG-2 is at least one of the upstream factors sufficient to bring downstream ESCRT III machinery via ALIX and ESCRT-I/II to the damaged endolysosomal membrane. Of course, it is likely that other signals, such as lysobisphosphatidic acid (LBPA) (41), Annexin A7 (17), phospho-Rab8A (42), could also contribute to increase the affinity and specificity of the process.

ESCRT-III machinery has been implicated in the plasma membrane repair. It has been shown that ESCRT-III machinery is recruited to the membrane damage site with the local influx of Ca^2+^ into the cytoplasm acting as the trigger for the recruitment of Ca^2+^ chelator proteins, possibly ALG-2 (11, 12). However, no one has recruited the entire membrane repair machinery in vitro. By drawing parallels between the plasma membrane repair and the endolysosomal repair, our reconstitution experiments elucidate the key mechanism for endolysosomal membrane repair.

Cell based studies on endolysosomal membrane repair have found that silencing of only ALIX or ESCRT-I/II pathway is insufficient to stall endolysosomal membrane repair (9, 10, 43). From our reconstitution experiments we explain this observation by demonstrating that ALG-2 can bind to both ALIX as well as ESCRT-I protein. This can subsequently lead to recruitment of ESCRT-III machinery from two parallel pathways. Therefore, our reconstitution experiments explain why silencing of ALIX or TSG 101 individually is insufficient to abrogate membrane repair and that silencing of only one gene (out of ALIX and TSG101) can be compensated by the alternate parallel pathway of ESCRT-III recruitment.

The prion-like propagation of molecular aggregates, in particular aggregated tau, is thought to be a prominent mechanism for the spread of misfolded tau in Alzheimer’s Disease (AD) and Frontotemporal Degeneration (FTD) (5). Endolysosomal escape in the receiver cell is a key step in the cell-to-cell spread of aggregated tau (44). A recent CRISPR interference screen-based study showed that knocking down ESCRT components, CHMP6 or CHMP2A together with CHMP2B increased endolysosomal membrane leakiness and promoted cytoplasmic entry of tau aggregates (6). Our observation that ALG-2 can recruit ESCRT-III machinery (starting at CHMP6) via the ESCRT-I/II pathway shows how CHMP6 is connected biochemically to the other main players in ESCRT-based lysosomal membrane repair.

## Materials and Methods

### Materials

The lipids 1,2-dioleoyl-sn-glycero-3-phosphocholine (DOPC) and 1,2-dioleoyl-sn-glycero-3-phospho-L-serine (sodium salt) (DOPS) were obtained from Avanti Polar Lipids (Alabaster, AL). Lipid fluorophore 1,2 – Dioleoyl-sn-glycero-3-phosphoethanolamine labeled with Atto 647 (Atto 647 DOPE) was purchased from Sigma Aldrich (St. Louis, MO). HEPES, NaCl, EGTA, and fatty acid free bovine albumin (BSA) were obtained from Fisher Scientific (Rochester, NY). All commercial reagents were used without further purification.

### GUV formation

GUVs containing DOPC (89.5, 69.5, or 49.5 mol%), DOPS (10, 30, or 50 mol%), and the lipid fluorophore Atto 647 DOPE (0.5 mol%) were prepared in 270 mOsm sucrose using PVA-gel hydration-based method as in Weinberger et al. (45). Briefly, lipids were mixed in chloroform at a total concentration of 1 mM. The 40 μL solution of the lipid mixture was spread on a 5% w/v PVA film dried on a 25 x 25 mm coverslip (VWR, Radnor, PA) and then put under vacuum for at least 2 h to form a dry lipid film. The dried lipid film was hydrated with 500 μL of 270 mOsm sucrose solution for 2 h at room temperature to produce GUV dispersion which was collected and stored in a 1.5 ml microcentrifuge tube.

### Protein purification

ALG-2 was purified based on the protocol described in McGourthy et al. (46). Briefly, N-terminal 6x His tagged ALG-2 was expressed in *E. coli* BL21 (DE3) cells in LB medium supplemented with kanamycin (50 μg/ml), induced at 0.8 OD with 0.5 mM IPTG at 37 °C for 3 h. After lysis through tip sonication in lysis buffer (50 mM Tris pH 7.4, 150 mM NaCl, 0.2 mM TCEP) the expressed protein was extracted from the supernatant using NiNTA resin (QIAGEN, Germantown, MD). The resulting eluate from the NiNTA resin using lysis buffer supplemented with 250 mM Imidazole pH 7.4, was loaded onto the Superdex 75 16/60 column (GE Healthcare) for gel filtration. Subsequently, the resulting solution was purified using anion exchange chromatography using 5 ml HiTrap Q HP (Cytiva, Marlborough, MA). Finally, the eluate was loaded on an equilibrated Superdex 75 16/60 column (GE Healthcare), and the protein purity assessed using sodium dodecyl sulfate polyacrylamide gel electrophoresis (SDS-PAGE). Concentration of the purified protein was calculated by measuring the absorbance at 280 nm. Finally, protein was concentrated to approximately 50 μM and stored at −80°C in small aliquots.

ESCRT-I (TSG101, VPS28, VPS37B, and MVB12A) and full-length ALIX were expressed in HEK293 cells. ESCRT-I had a Strep-tagged VPS28 subunit and ALIX was C-terminally strep-tagged, and were initially purified on StrepTactin Sepharose (IBA), followed by gel filtration chromatography on a Superdex 200 16/60 column (GE Healthcare), in 50 mM Tris pH 7.4, 300 mM NaCl, 0.1 mM TCEP.

ESCRT-II (EAP45, EAP30, EAP20) was expressed in E. coli Rosetta2 (DE3) at 20°C overnight in LB medium, with an N-terminal TEV-cleavable 6x His tag and purified on NiNTA resin, followed by gel filtration chromatography on a Superdex 200 16/60 column (GE Healthcare), in 50 mM Tris pH 7.4, 300 mM NaCl, 0.1 mM TCEP. The 6x His tag was removed by TEV protease digestion followed by passage over NiNTA followed by a final gel filtration chromatography on a Superdex 200 16/60 column (GE Healthcare). The final purified protein was concentrated to approximately 10 μM and snap frozen on liquid nitrogen in small aliquots.

ESCRT-III (CHMP6, CHMP4B, CHMP2A, CHMP3) proteins were purified as described in Carlson et al. (18). Briefly, they were expressed individually as an N-terminal TEV-cleavable 6x His-MBP fusion in E. coli Rosetta2 (DE3) at 20°C overnight in LB medium. Cells were lysed by tip sonication, and initial purification was carried out on NiNTA resin. For the case of CHMP4B, the 6x His-MBP fusion was further purified on a Superdex 200 16/60 column (GE Healthcare), in 50 mM Tris pH 7.4, 100 mM NaCl, 0.1 mM TCEP. The 6x His-MBP tag was removed by TEV protease digestion at low micromolar concentrations, followed by gel filtration chromatography on a Superdex 75 16/60 column (GE Healthcare), in 50 mM Tris pH 7.4, 100 mM NaCl, 0.1 mM TCEP. CHMP4B eluted as a monodisperse sample, typically with a concentration of approximately 400 nM. Unconcentrated CHMP4B fractions were kept at 4 °C and used within 72 h of purification because CHMP4B formed soluble aggregates over time and freeze-thaw resulted in a loss of material. The other ESCRT-III subunits were purified analogous to CHMP4B, except that the final proteins had higher concentration (approximately 20-50 μM) and could be snap frozen on liquid nitrogen without aggregation or loss of material. 6x His tagged VPS4B was purified similar to ESCRT-III proteins except an anion exchange step was added between the metal affinity (NiNTA) purification and the final gel filtration chromatography step.

Fluorophore labeling was performed overnight at 4°C using cysteine reactive dyes on engineered N-terminal cysteines (ESCRT-III, VPS4B), or A78C for ALG-2, or on native surface-exposed cysteines (ALIX). Specifically, Atto 488 maleimide (Sigma-Aldrich) was used for labeling ALG-2 and CHMP4B; sulpho-Cy3 maleimide (Fischer Scientific, Hampton, NH) for labelling ALIX and CHMP3, and Lumidyne 655 maleimide (Lumidyne Technologies, NY) for labelling VPS4B. Labeling was performed on the fusion proteins before TEV digest for proteins with a TEV cleavage site. Excess dye was removed by passing the protein-dye mixture through two PD10 columns (Cytiva, Marlborough, MA) sequentially. Final step in every protein purification was gel filtration chromatography, so that the monodisperse state of the (labelled) protein could be ensured. Labeling efficiencies were normally 50–100%, except for ALG-2 which had a 25% labeling efficiency.

### Reconstitution Reactions and Confocal Microscopy

The incubation reactions were set up in a microcentrifuge tube at room temperature before transferring to a Lab-Tek II chambered cover glass (Fisher Scientific) for imaging. The imaging chamber was pre-coated with a 5 mg/ml solution of fatty acid free Bovine Serum Albumin for 30 min and washed three times with the reaction buffer (25 mM HEPES at pH 7.4, 125 mM NaCl, and 0.2 mM TCEP, 280 mOsm) before transferring the reactants from the microcentrifuge tubes. 15 μL of GUVs were mixed with 120 μL of reaction buffer containing proteins at concentrations stated in ***Results***. After 15-min incubation images were acquired on a Nikon A1 confocal microscope with a 63× Plan Apochromat 1.4 numerical aperture (NA) objective. Three replicates were performed for each experimental condition. Identical laser power and gain settings were used for each set of replicates.

### Immunofluorescence

U2OS osteosarcoma cells were maintained at 37°C under 5% CO_2_ and propagated using Dulbeco’s modified Eagle’s medium (DMEM) (no. 11965-084, Gibco) supplemented with 10% v/v heat-inactivated fetal bovine serum (FBS). Prior to experimentation, cells were seeded into 35 mm glass bottom dishes (no. P35G-1.5-14-C, MatTek) and treated with 1 mM L-Leucyl-L-Leucine methyl ester (hydrochloride) (LLOME) (no. L7393, Sigma-Aldrich) dissolved in dimethtyl sulfoxide (DMSO), or DMSO vehicle control for 15 minutes. Cells were then fixed in 4% paraformaldehyde (Electron Microscopy Sciences) in PBS for 15 minutes at room temperature. Cells were then rinsed once with PBS prior to being permeabilized with 0.02% Digitonin (no. BN2006, Thermo Fisher) in PBS for 10 minutes at room temperature.

Cells were then rinsed once with PBS and blocked with 2% w/v Bovine serum albumin (BSA) in PBS for 30 minutes at room temperature. Blocked cells were rinsed three times with PBS prior to incubation with primary antibodies diluted in 2% BSA in PBS. ALG-2 was detected using the PDCD6 Rabbit Polyclonal Antibody (no. 12303-1-AP, Thomas Scientific), ALIX, using the PDCD6IP Mouse Monoclonal Antibody (no. MA1-83977, Thermo Fisher), TSG101, using the primary-conjugated Mouse Monoclonal Antibody (no. sc-7964 AF647, Santa Cruz Biotechnology), and Gal3, using the primary-conjugated Mouse Monoclonal Antibody (no. sc-32790 AF594, Santa Cruz Biotechnology). ALG-2 and ALIX antibodies were diluted 1:200 with 2% BSA in PBS and incubated with cells for 1 hour at room temperature.

To remove non-specific antibody binding, cells were washed three times with PBS and antibody binding was probed using secondary-conjugated Goat antibodies targeting the species of ALG-2 and ALIX antibodies (no. ab175652, ab150113, abcam). Secondary antibodies were diluted 1:500 with 2% BSA in PBS and incubated with cells in the dark for 30 minutes at room temperature. Cells were washed an additional three times in PBS to remove non-specific secondary antibodies and incubated with primary-conjugated antibodies against TSG101 and Gal3 diluted 1:200 in 2% BSA in PBS for 30 minutes in the dark at room temperature. Cells were washed a final three times in PBS and imaged immediately.

### Image Analysis

Puncta recognition analysis was performed by custom-made scripts in Python. The lipid channel was used to identify GUVs using Yolov5 developed by Ultralytics (https://github.com/ultralytics/yolov5.git). For each recognized GUV, the corresponding protein channel images were pre-processed by using *threshold_li* (from scikit-image) over the dataset on the same experiment to choose a value for performing background subtraction. *Fast Nl Means Denoising*, *Gaussian Blur*, and *Equalize Histogram* (all from OpenCV i.e., cv2) functions were performed after background subtraction. Subsequently a binary mask was created using threshold_otsu (from scikit-image). The foreground regions with areas larger than 5 pixels were counted as a puncta. Finally, the colocalization information was then extracted by comparing the punctum position across the protein channels by proximity criterion defined such that two puncta with centroids closer than 0.25 times the diameter of the GUV were considered colocalized.

ImageJ was used for the image analysis of immunofluorescence data. U2OS cells stained via immunofluorescence after treatment with DMSO or LLOME were subject to coincident area quantification. Images of cells were thresholded equally to select regions of interest containing all fluorescent markers. The average pixel area of the intersecting channels was calculated and compared between DMSO and LLOME treated samples.

### Statistical analysis

Statistical analysis was performed with GraphPad Prism 9.0 (La Jolla, CA, USA). The data of GUV binding assay was analyzed by student’s t-test and one-way ANOVA. Significance between the two calculated areas was determined using a student’s two-tailed unpaired t test. *P* < 0.05 was considered statistically significant.

## Acknowledgements

We thank A. King Cada for generating Fig. 7, members of the Hurley lab for helpful discussions and critical feedback on the manuscript, and the M. Rape lab for the ALG-2 expression construct.

## References

1. K. P. Bohannon, P. I. Hanson, ESCRT puts its thumb on the nanoscale: Fixing tiny holes in endolysosomes. Curr Opin Cell Biol 65, 122–130 (2020).

2. J. Staring, M. Raaben, T. R. Brummelkamp, Viral escape from endosomes and host detection at a glance. J Cell Sci 131 (2018).

3. M. van Hees et al., New approaches to moderate CRISPR-Cas9 activity: Addressing issues of cellular uptake and endosomal escape. Mol Ther 30, 32–46 (2022).

4. Z. Wu, T. Li, Nanoparticle-Mediated Cytoplasmic Delivery of Messenger RNA Vaccines: Challenges and Future Perspectives. Pharm Res 38, 473–478 (2021).

5. J. Vaquer-Alicea, M. I. Diamond, Propagation of Protein Aggregation in Neurodegenerative Diseases. Annual Review of Biochemistry 88, 785–810 (2019).

6. J. J. Chen et al., Compromised function of the ESCRT pathway promotes endolysosomal escape of tau seeds and propagation of tau aggregation. J Biol Chem 294, 18952–18966 (2019).

7. J. H. Hurley, ESCRTs are everywhere. EMBO Journal 34, 2398–2407 (2015).

8. M. Vietri, M. Radulovic, H. Stenmark, The many functions of ESCRTs. Nat Rev Mol Cell Biol 21, 25–42 (2020).

9. M. L. Skowyra, P. H. Schlesinger, T. V. Naismith, P. I. Hanson, Triggered recruitment of ESCRT machinery promotes endolysosomal repair. Science 360 (2018).

10. M. Radulovic et al., ESCRT-mediated lysosome repair precedes lysophagy and promotes cell survival. EMBO J 37 (2018).

11. A. J. Jimenez et al., ESCRT machinery is required for plasma membrane repair. Science 343, 1247136 (2014).

12. L. L. Scheffer et al., Mechanism of Ca(2)(+)-triggered ESCRT assembly and regulation of cell membrane repair. Nat Commun 5, 5646 (2014).

13. G. E. Kass, S. Orrenius, Calcium signaling and cytotoxicity. Environmental Health Perspectives 107, 25–35 (1999).

14. K. A. Christensen, J. T. Myers, J. A. Swanson, pH-dependent regulation of lysosomal calcium in macrophages. Journal of Cell Science 115, 599–607 (2002).

15. K. W.-H. Lo, Q. Zhang, M. Li, M. Zhang, Apoptosis-Linked Gene Product ALG-2 Is a New Member of the Calpain Small Subunit Subfamily of Ca2+-Binding Proteins. Biochemistry 38, 7498–7508 (1999).

16. M. Maki, H. Suzuki, H. Shibata, Structure and function of ALG-2, a penta-EF-hand calcium-dependent adaptor protein. Sci China Life Sci 54, 770–779 (2011).

17. S. L. Sonder et al., Annexin A7 is required for ESCRT III-mediated plasma membrane repair. Sci Rep 9, 6726 (2019).

18. L. A. Carlson, J. H. Hurley, In vitro reconstitution of the ordered assembly of the endosomal sorting complex required for transport at membrane-bound HIV-1 Gag clusters. Proc Natl Acad Sci U S A 109, 16928–16933 (2012).

19. R. Pires et al., A Crescent-Shaped ALIX Dimer Targets ESCRT-III CHMP4 Filaments. Structure 17, 843–856 (2009).

20. Q. Zhai et al., Activation of the Retroviral Budding Factor ALIX. Journal of Virology 85, 9222–9226 (2011).

21. S. Sun et al., ALG-2 activates the MVB sorting function of ALIX through relieving its intramolecular interaction. Cell Discov 1, 15018 (2015).

22. M. Missotten, A. Nichols, K. Rieger, R. Sadoul, Alix, a novel mouse protein undergoing calcium-dependent interaction with the apoptosis-linked-gene 2 (ALG-2) protein. Cell Death & Differentiation 6, 124–129 (1999).

23. M. Bajorek et al., Structural basis for ESCRT-III protein autoinhibition. Nature Structural & Molecular Biology 16, 754–762 (2009).

24. S. Lata et al., Structural Basis for Autoinhibition of ESCRT-III CHMP3. Journal of Molecular Biology 378, 818–827 (2008).

25. S. Shim, L. A. Kimpler, P. I. Hanson, Structure/Function Analysis of Four Core ESCRT-III Proteins Reveals Common Regulatory Role for Extreme C-Terminal Domain. Traffic 8, 1068–1079 (2007).

26. S. Tang et al., Structural basis for activation, assembly and membrane binding of ESCRT-III Snf7 filaments. eLife 4 (2015).

27. J. Schöneberg et al., ATP-dependent force generation and membrane scission by ESCRT-III and Vps4. Science 362, 1423–1428 (2018).

28. M. Babst, D. J. Katzmann, E. J. Estepa-Sabal, T. Meerloo, S. D. Emr, Escrt-III. Developmental Cell 3, 271–282 (2002).

29. B. Yang, G. Stjepanovic, Q. Shen, A. Martin, J. H. Hurley, Vps4 disassembles an ESCRT-III filament by global unfolding and processive translocation. Nature Structural & Molecular Biology 22, 492–498 (2015).

30. T. Obita et al., Structural basis for selective recognition of ESCRT-III by the AAA ATPase Vps4. Nature 449, 735–739 (2007).

31. M. D. Stuchell-Brereton et al., ESCRT-III recognition by VPS4 ATPases. Nature 449, 740–744 (2007).

32. K. Katoh et al., The penta-EF-hand protein ALG-2 interacts directly with the ESCRT-I component TSG101, and Ca2+-dependently co-localizes to aberrant endosomes with dominant-negative AAA ATPase SKD1/Vps4B. Biochemical Journal 391, 677–685 (2005).

33. C. Langelier et al., Human ESCRT-II Complex and Its Role in Human Immunodeficiency Virus Type 1 Release. Journal of Virology 80, 9465–9480 (2006).

34. A. Hierro et al., Structure of the ESCRT-II endosomal trafficking complex. Nature 431, 221–225 (2004).

35. H. Teo, O. Perisic, B. González, R. L. Williams, ESCRT-II, an Endosome-Associated Complex Required for Protein Sorting. Developmental Cell 7, 559–569 (2004).

36. Y. J. Im, J. H. Hurley, Integrated Structural Model and Membrane Targeting Mechanism of the Human ESCRT-II Complex. Developmental Cell 14, 902–913 (2008).

37. C. Yorikawa et al., Human CHMP6, a myristoylated ESCRT-III protein, interacts directly with an ESCRT-II component EAP20 and regulates endosomal cargo sorting. Biochemical Journal 387, 17–26 (2005).

38. D. L. Thiele, P. E. Lipsky, Mechanism of L-leucyl-L-leucine methyl ester-mediated killing of cytotoxic lymphocytes: dependence on a lysosomal thiol protease, dipeptidyl peptidase I, that is enriched in these cells. Proceedings of the National Academy of Sciences 87, 83–87 (1990).

39. S. Aits et al., Sensitive detection of lysosomal membrane permeabilization by lysosomal galectin puncta assay. Autophagy 11, 1408–1424 (2015).

40. J. Westman et al., Calcium-dependent ESCRT recruitment and lysosome exocytosis maintain epithelial integrity during Candida albicans invasion. Cell Rep 38, 110187 (2022).

41. C. Bissig et al., Viral infection controlled by a calcium-dependent lipid-binding module in ALIX. Dev Cell 25, 364–373 (2013).

42. S. Herbst et al., LRRK2 activation controls the repair of damaged endomembranes in macrophages. EMBO J 39, e104494 (2020).

43. J. Jia et al., Galectin-3 Coordinates a Cellular System for Lysosomal Repair and Removal. Dev Cell 52, 69–87 e68 (2020).

44. B. Frost, R. L. Jacks, M. I. Diamond, Propagation of Tau Misfolding from the Outside to the Inside of a Cell. Journal of Biological Chemistry 284, 12845–12852 (2009).

45. A. Weinberger et al., Gel-Assisted Formation of Giant Unilamellar Vesicles. Biophysical Journal 105, 154–164 (2013).

46. C. A. McGourty et al., Regulation of the CUL3 Ubiquitin Ligase by a Calcium-Dependent Co-adaptor. Cell 167, 525–538 e514 (2016).

